# Genome-wide genetic data on ~500,000 UK Biobank participants

**DOI:** 10.1101/166298

**Authors:** Clare Bycroft, Colin Freeman, Desislava Petkova, Gavin Band, Lloyd T. Elliott, Kevin Sharp, Allan Motyer, Damjan Vukcevic, Olivier Delaneau, Jared O’Connell, Adrian Cortes, Samantha Welsh, Gil McVean, Stephen Leslie, Peter Donnelly, Jonathan Marchini

**Affiliations:** Wellcome Trust Center for Human Genetics, University of Oxford, UK; Department of Statistics, University of Oxford, UK; Centre for Systems Genomics and the Schools of Mathematics and Statistics, and BioSciences, The University of Melbourne, Parkville, Victoria, Australia; Murdoch Children’s Research Institute, Parkville, Victoria, Australia; Department of Genetic Medicine and Development, University of Geneva, 1 Michel Servet, Geneva, CH1211, Switzerland; Swiss Institute of Bioinformatics, University of Geneva, 1 Michel Servet, Geneva, CH1211, Switzerland; Institute of Genetics and Genomics in Geneva, University of Geneva, 1 Michel Servet, Geneva, CH1211, Switzerland; Illumina Ltd, Chesterford Research Park, Little Chesterford, Essex, CB10 1XL, United Kingdom; Nuffield Department of Clinical Neurosciences, Division of Clinical Neurology, John Radcliffe Hospital, University of Oxford, Oxford OX3 9DU, United Kingdom; UK Biobank, Units 1-4 Spectrum Way, Adswood, Stockport, Cheshire, SK3 0SA, UK; Big Data Institute, Li Ka Shing Centre for Health Information and Discovery, University of Oxford, Oxford OX3 7LF, United Kingdom

**Keywords:** UK Biobank, Genotypes, Quality control, Population structure, Relatedness, Phasing, Imputation, HLA Imputation, GWAS, PheWAS

## Abstract

The UK Biobank project is a large prospective cohort study of ~500,000 individuals from across the United Kingdom, aged between 40-69 at recruitment. A rich variety of phenotypic and health-related information is available on each participant, making the resource unprecedented in its size and scope. Here we describe the genome-wide genotype data (~805,000 markers) collected on all individuals in the cohort and its quality control procedures. Genotype data on this scale offers novel opportunities for assessing quality issues, although the wide range of ancestries of the individuals in the cohort also creates particular challenges. We also conducted a set of analyses that reveal properties of the genetic data – such as population structure and relatedness – that can be important for downstream analyses. In addition, we phased and imputed genotypes into the dataset, using computationally efficient methods combined with the Haplotype Reference Consortium (HRC) and UK10K haplotype resource. This increases the number of testable variants by over 100-fold to ~96 million variants. We also imputed classical allelic variation at 11 human leukocyte antigen (HLA) genes, and as a quality control check of this imputation, we replicate signals of known associations between HLA alleles and many common diseases. We describe tools that allow efficient genome-wide association studies (GWAS) of multiple traits and fast phenome-wide association studies (PheWAS), which work together with a new compressed file format that has been used to distribute the dataset. As a further check of the genotyped and imputed datasets, we performed a test-case genome-wide association scan on a well-studied human trait, standing height.

## 1 Introduction

The UK Biobank project is a large prospective cohort study of ~500,000 individuals from across the United Kingdom, aged between 40-69 at recruitment [1]. A rich variety of phenotypic and health-related information is available on each participant, making the resource unprecedented in its size and scope. The data contains self reported information, including basic demographics, diet, and exercise habits; extensive physical and cognitive measurements; with other sources of health-related information such as medical records and cancer registers being integrated and followed up over the course of the participants’ lives [2]. The baseline information has, and will be, extended in a number of ways [3]. For example, many blood and urine biomarkers are being measured; and medical imaging of brain [4], heart, bones, carotid arteries and abdominal fat is being carried out on a large subset (~100,000) of participants [5].

Understanding the role that genetics plays in phenotypic and disease variation, and its potential interactions with other factors, provides a critical route to a better understanding of human biology. It is anticipated that this will lead to more successful drug development [6], and potentially to more efficient and personalised treatments and to better diagnoses. As such, a key component of the UK Biobank resource has been the collection of genome-wide genetic data on every participant using a purpose-designed genotyping array [7]. An interim release of genotype data on ~150,000 UK Biobank participants (May 2015) [8] has already facilitated numerous studies [9]. These exploit the UK Biobank’s substantial sample size, extensive phenotype information, and genome-wide genetic information to study the often subtle and complex effects of genetics on human traits and disease, and its potential interactions with other factors [10-15].

In this paper we describe the genetic dataset on the full ~500,000 participants, together with a range of quality control procedures, which have been undertaken on the genotype data in the hope of facilitating its wider use. To achieve this we designed and implemented a quality control (QC) pipeline that addresses challenges specific to the experimental design, scale, and diversity of this dataset. Raw data from the genotyping experiments will be available from UK Biobank. We also conducted a set of analyses that reveal properties of the genetic data – such as population structure and relatedness – that can be important for downstream analyses. In addition, we phased and imputed genotypes into the dataset, using computationally efficient methods combined with the Haplotype Reference Consortium (HRC) [16] and UK10K haplotype resources [17]. This increases the number of testable variants by over 100-fold to ~96 million variants. We also imputed classical allelic variation at 11 human leukocyte antigen (HLA) genes, and as a QC check of this imputation, we replicate signals of known associations between HLA alleles and many common diseases. We describe tools that allow efficient genome-wide association studies (GWAS) of multiple traits and fast phenome-wide association studies (PheWAS), which work together with a new compressed file format that has been used to distribute the dataset. As a further check of the genotyped and imputed datasets, we performed a test-case genome-wide association scan on a well-studied human trait, standing height.

## 2 Results

### 2.1 High quality genotype calling on novel array

#### 2.1.1 Purpose-designed genotyping array

The data release contains genotypes of 488,377 UK Biobank participants. These were assayed using two very similar genotyping arrays. A subset of 49,950 participants involved in the UK Biobank Lung Exome Variant Evaluation (UK BiLEVE) study were genotyped using the Applied Biosystems™ UK BiLEVE Axiom™ Array by Affymetrix^1^ (807,411 markers), which is described elsewhere [15]. Following this, 438,427 participants were genotyped using the closely-related Applied Biosystems™ UK Biobank Axiom™ Array (825,927 markers). Both arrays were purpose-designed specifically for the UK Biobank genotyping project and share 95% of marker content [7]. The marker content of the UK Biobank Axiom™ array was chosen to capture genome-wide genetic variation (single nucleotide polymorphism (SNPs) and short insertions and deletions (indels)), and is summarised in **Figure 1**. Many markers were included because of known associations with, or possible roles in, phenotypic variation, particularly disease. A notable example is the inclusion of two variants, rs429358 and rs7412, which define the isoforms of the apolipoprotein E (*APoE*) gene known to be associated with risk of Alzheimer’s disease [7] and other conditions. Neither marker is easy to type using array technologies; as a consequence of this they have not always been assayed on earlier arrays. The array also includes coding variants across a range of minor allele frequencies (MAFs), including rare markers (<1% MAF); and markers that provide good genome-wide coverage for imputation in European populations in the common (>5%) and low frequency (1-5%) MAF ranges. Further details of the array design are in [7].

**Figure 1.**
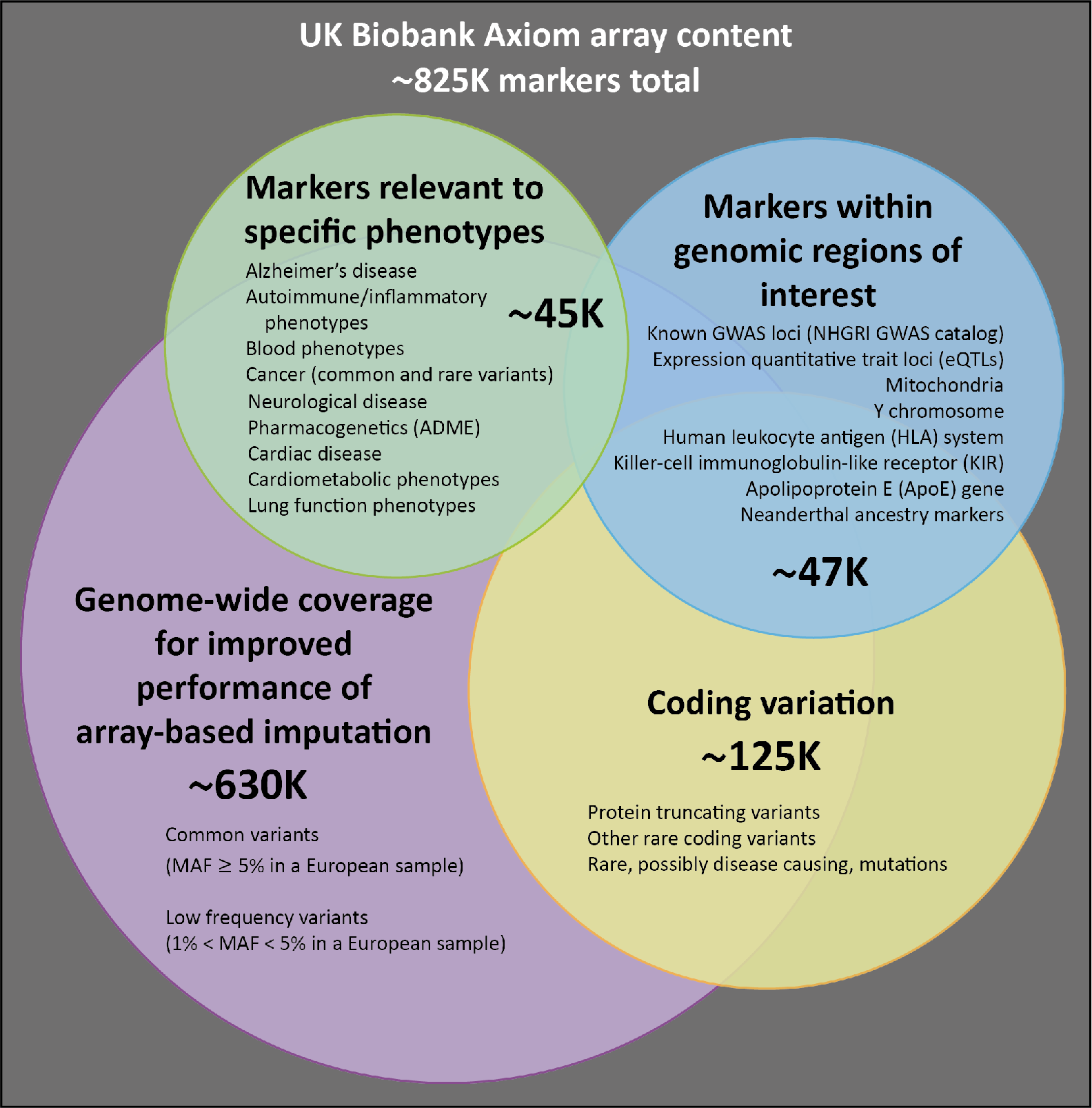
Summary of UK Biobank genotyping array content. This is a schematic representation of the different categories of content on the UK Biobank Axiom array. Numbers indicate the approximate count of markers within each category, ignoring any overlap. A more detailed description of the array content is available in [7].

#### 2.1.2 DNA extraction and genotype calling

Blood samples were collected from participants on their visit to a UK Biobank assessment centre and the samples are stored at the UK Biobank facility in Stockport, UK [18]. Over a period of 18 months (Nov. 2013 – Apr. 2015) samples were retrieved, DNA was extracted, and 96-well plates of 94 50μl aliquots were shipped to Affymetrix Research Services Laboratory for genotyping. Special attention was paid in the automated sample retrieval process at UK Biobank to ensure that experimental units such as plates or timing of extraction did not correlate systematically with baseline phenotypes such as age, sex, and ethnic background, or the time and location of sample collection. Full details of the UK Biobank sample retrieval and DNA extraction process are described in [19, 20].

On receipt of DNA samples, Affymetrix processed samples on the GeneTitan® Multi-Channel (MC) Instrument in 96-well plates containing 94 UK Biobank samples and two control samples from the 1000 Genomes Project [21]. Genotypes were then called from the array intensity data, in units called “batches” which consist of multiple plates. Across the entire cohort, there were 106 batches of ~4,700 UK Biobank samples each (**Supplementary Material, Table S11**). Following the earlier interim data release, Affymetrix developed a custom genotype calling pipeline that is optimized for biobank-scale genotyping experiments, which takes advantage of the multiple-batch design [22]. This pipeline was applied to all samples, including the~150,000 samples that were part of the interim data release. Consequently, some of the genotype calls for these samples may differ between the interim data release and this final data release (see Section 2.1.6).

Routine quality checks were carried out during the process of sample retrieval, DNA extraction [19], and genotype calling [23]. Any sample that did not pass these checks was excluded from the resulting genotype calls. The custom-designed arrays contain a number of markers that had not been previously typed using Affymetrix genotype array technology. As such, Affymetrix also applied a series of checks to determine whether the genotyping assay for a given marker was successful, either within a single batch, or across all samples. Where these newly-attempted assays were not successful, Affymetrix excluded the markers from the data delivery (see **Supplementary Material** for details). This resulted in a set of genotype calls for 489,212 samples at 812,428 unique markers (bi-allelic SNPs and Indels) from both arrays, with which we conducted further quality control and analysis (**Table 1**).

**Table 1.** The number of markers and samples by genotyping array at main stages of the UK Biobank genotyping experiment. “Data delivery from Affymetrix” refers to the data produced by Affymetrix after applying their filtering (see Supplementary Material). “Released data” refers to the released genotype data, after applying QC as described in Sections 2.1.4 and 2.1.5.

|  |  | UK BiLEVE<br>Axiom array<br>only | UK Biobank<br>Axiom array<br>only | Both arrays | Total |
| --- | --- | --- | --- | --- | --- |
| Included in experiment | Number of samples sent to Affymetrix (including duplicates) | 50561 | 443568 | 0 | 494078 |
| Included in data delivery from Affymetrix | Number of markers | 18019 | 34313 | 760096 | 812428 |
|  | Number of samples (including duplicates) | 50520 | 438692 | 0 | 489212 |
| Included in released data | Number of markers | 17536 | 34197 | 753693 | 805426 |
|  | Number of unique samples | 49950 | 438427 | 0 | 488377 |

#### 2.1.3 Quality control in a large-scale, ethnically diverse cohort

Our QC pipeline was designed specifically to accommodate the large-scale dataset of ethnically diverse participants, genotyped in many batches (106), using two slightly different novel arrays, and which will be used by many researchers to tackle a wide variety of research questions. Participants reported their ethnic background by selecting from a fixed set of categories [24]. While the majority (94%) of individuals report their ethnic background as within the broad-level group “White”, there are still ~22,000 individuals with a self-reported ethnic background originating outside Europe (**Table 2**). This ethnic diversity implies genetic diversity, which we observe directly in the genotypes as allele frequency differences, and has implications for QC. Some commonly used QC tests are ineffective in the context of strong population structure if applied without taking this into account. For example, testing for departures from Hardy-Weinberg equilibrium (HWE) is a common approach for identifying markers that have been genotyped poorly [25-27], but departures from HWE can be expected in the context of population structure because of differences in allele frequencies across populations. We used approaches based on principal component analysis (PCA) to account for population structure in both marker and sample-based QC.

**Table 2.** Proportions of self-reported ethnic groups among 488,377 genotyped UK Biobank participants. Categories of self-reported ethnic background (UK Biobank data field 21000) and broader-level ethnic groups are shown here to reflect the two-layer branching structure of the ethnic background section in the UK Biobank touchscreen questionnaire [24]. Participants first picked one of the broader-level ethnic groups (e.g White), and were then prompted to select one of the categories within that group (e.g Irish). The broader-level groups are also shown here as an ethnic background category (e.g “White” in column two) because a small proportion of participants only responded to the first question. In this table we also combine the category “Other ethnic group” with an aggregated non-response category “Not stated”, which includes all participants who did not know their ethnic group, or stated that they preferred not answer, or did not answer the first question.

| Ethnic group | Self-reported ethnic background | Percentage of genotyped UK Biobank participants |
| --- | --- | --- |
| **White** |  | 94.23 |
|  | British | 88.26 |
|  | Any other white background | 3.24 |
|  | Irish | 2.61 |
|  | White | 0.11 |
| **Asian or Asian British** |  | 1.94 |
|  | Indian | 1.17 |
|  | Pakistani | 0.36 |
|  | Any other Asian background | 0.36 |
|  | Bangladeshi | 0.05 |
|  | Asian or Asian British | 0.01 |
| **Black or Black British** |  | 1.57 |
|  | Caribbean | 0.88 |
|  | African | 0.66 |
|  | Any other Black background | 0.02 |
|  | Black or Black British | 0.01 |
| **Chinese** |  | 0.31 |
|  | Chinese | 0.31 |
| **Mixed** |  | 0.58 |
|  | Any other mixed background | 0.2 |
|  | White and Asian | 0.16 |
|  | White and Black Caribbean | 0.12 |
|  | White and Black African | 0.08 |
|  | Mixed | 0.01 |
| **Other/Unknown** |  | 1.38 |
|  | Other ethnic group | 0.89 |
|  | Not stated | 0.48 |

#### 2.1.4 Marker-based QC

We identified poor quality markers using statistical tests designed primarily to check for consistency across experimental factors. Specifically we tested for batch effects, plate effects, departures from HWE, sex effects, array effects, and discordance across control replicates. See **Supplementary Material** for the details of each test, and **Figure S3** for examples of affected markers. For markers that failed at least one test in a given batch, we set the genotype calls in that batch to missing. We also provide a flag in the data release that indicates if the calls for a marker have been set to missing in a given batch. If there was evidence that a marker was not reliable across all batches, we excluded the marker from the data altogether. In order to attenuate population structure effects we applied all marker-based QC tests using a subset of 463,844 individuals with estimated European ancestry. We identified these individuals from the genotype data prior to conducting any QC by projecting all the UK Biobank samples on to the two major principal components of four 1000 Genomes populations (CEU, YRI, CHB and JPT) [28]. We then selected samples with principal component (PC) scores falling in the neighbourhood of the CEU cluster (see **Supplementary Material**).

Most QC metrics require a threshold beyond which to consider a marker ‘not reliable’. We used thresholds such that only strongly deviating markers would fail QC tests (see **Supplementary Material**), therefore allowing researchers to further refine the QC in whichever way is most appropriate for their study requirements. **Table 3** summarises the amount of data affected by applying these tests.

**Table 3.**
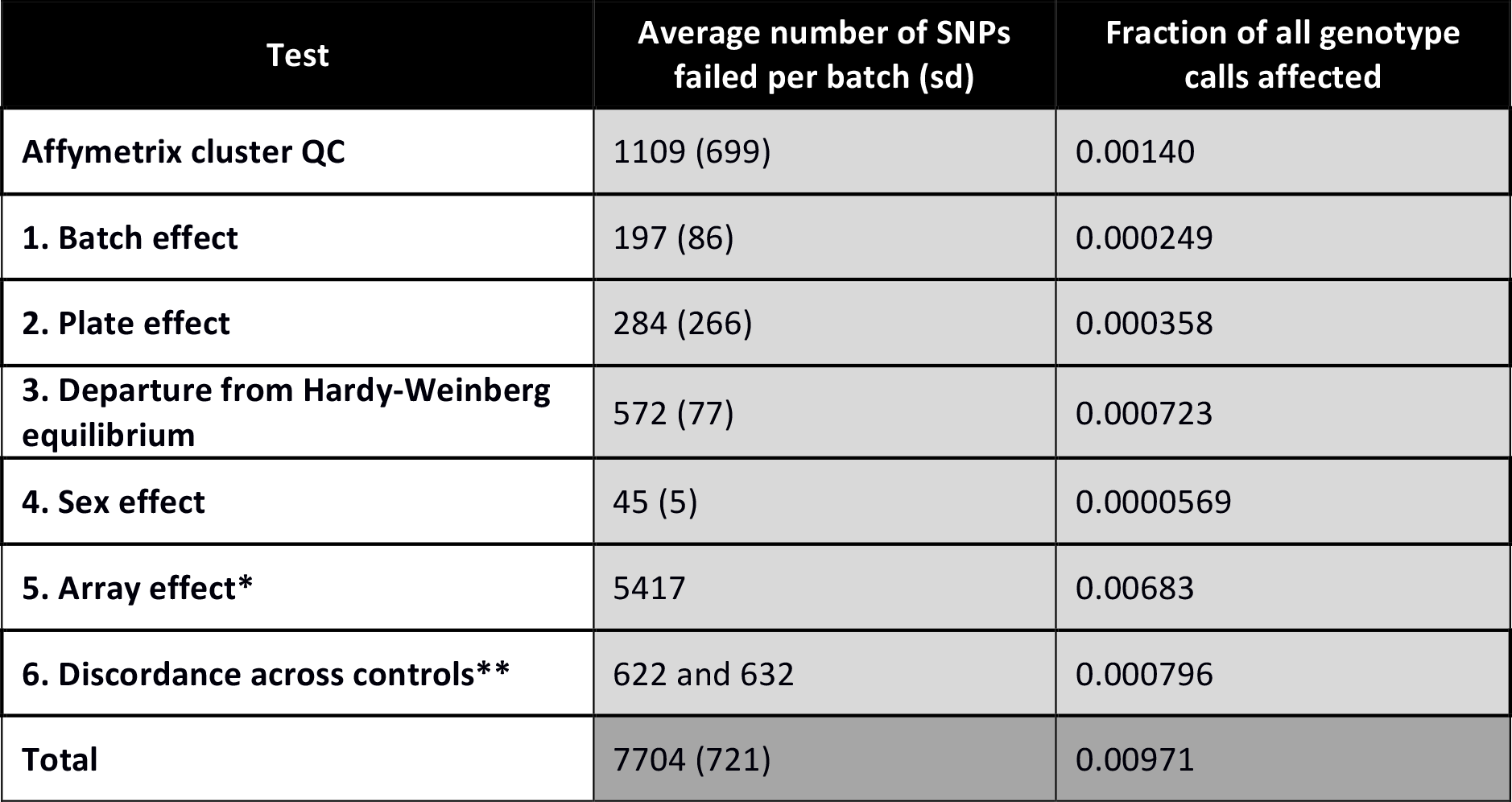
Failure rates for six marker-based quality tests. For all numbered tests a marker (or marker within a batch) was set to missing if the test yielded a p-value < 10^-12^, except in the case of Test 6, for which a marker was set to missing if the test yielded < 95% concordance. See Supplementary Material for details of each test. The total is not equal to the sum of all tests because it is possible for a marker to fail more than one test. Since the two arrays contain slightly different sets of markers, the total number of genotype calls used to compute the fractions is, *N_ukbb_ L_ukbb_* + *N_ukbl_ L_ukbl_*, where *N* and *L* refer to the numbers of markers and samples typed on the UK Biobank Axiom array (*ukbb*) and samples typed on the UK BiLEVE Axiom array (*ukbl*) within the Affymetrix data delivery (see Table S1). *The array effect test was applied across all batches and only for markers present on both arrays, so we simply report the total number of markers that failed this test. **The discordance test was applied across all batches, but not all markers are present on both arrays. The first value is the number of unique markers on the UK BiLEVE Axiom array that failed this test, and the second is for markers on the UK Biobank Axiom array.

#### 2.1.5 Sample-based QC

We identified poor quality samples using the metrics of missing rate and heterozygosity computed using a set of 605,876 high quality autosomal markers that were typed on both arrays (see **Supplementary Material** for criteria). Extreme values in one or both of these metrics can be indicators of poor sample quality due to, for example, DNA contamination [26]. The heterozygosity of a sample – the fraction of non-missing markers that are called heterozygous – can also be sensitive to natural phenomena, including population structure, recent admixture and parental consanguinity. We took extra measures to avoid misclassifying good quality samples because of these effects. For example, we adjusted heterozygosity for population structure by fitting a linear regression model with the first six principal components (PCs) in a PCA as predictors (**Figure 2A**). Using this adjustment we identified 968 samples with unusually high heterozygosity or >5% missing rate (**Supplementary Material**). A list of these samples is provided as part of the data release.

**Figure 2.**
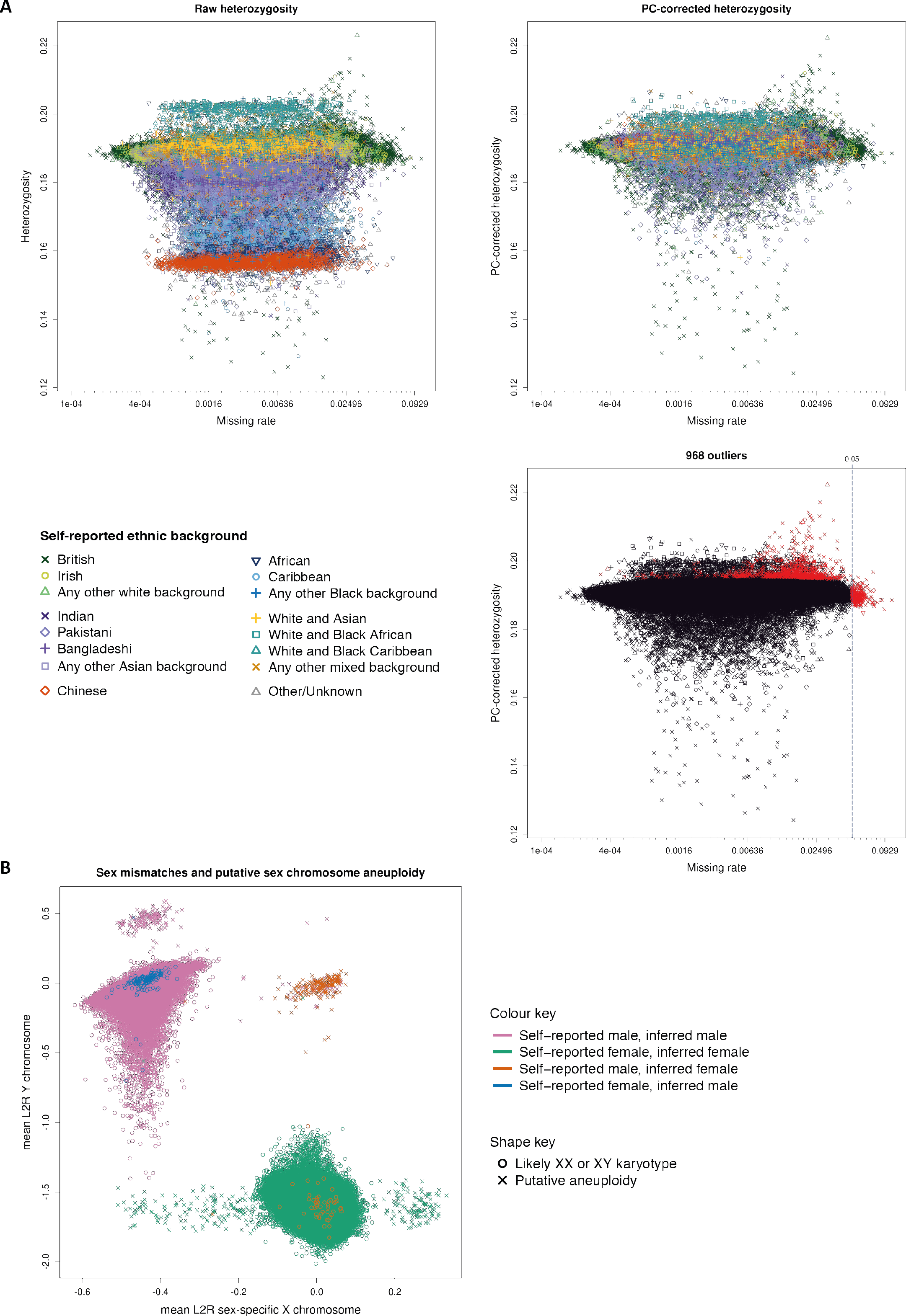
Summary of sample-based QC. (A) The three plots show heterozygosity and missing rates, which we used to flag poor quality samples. The two top plots show heterozygosity for each sample before (left) and after (right) correcting for ancestral background using PCs. The symbols (shapes and colours) indicate the self-reported ethnic background of each participant, as annotated in the legend. The third plot shows the set of samples we flagged as outliers (in red), and all other samples (in black), with shapes the same as for the other two plots. The vertical line shows the threshold we used to call samples as outliers on missing rate. In all plots missing rate data is transformed to the logit scale, but with the axis annotated with the original values. **(B)** Mean Log2 ratios (L2R) on X and Y chromosomes for each sample, indicating likely sex chromosome aneuploidy (see **Supplementary Material**). Samples which are most likely XX or XY are depicted with a circle. Other samples, which represent possible instances of sex chromosome aneuploidy, are indicated by a cross. There are 652 such samples in total; see **Supplementary Table S1** for counts. The colours of each symbol relate to different combinations of self-reported sex, and sex inferred by Affymetrix (from the genetic data), as indicated by the key. For almost all samples (99.9%) the self-reported and inferred sex are the same, but for a small number of samples (378) they do not match (see **Supplementary Material** for discussion).

We also conducted quality control specific to the sex chromosomes using a set of 15,766 high quality markers on the X and Y chromosomes. Affymetrix infers the sex of each individual based on the relative intensity of markers on the Y and X chromosomes [29]. Sex is also reported by participants, and mismatches between these sources can be used as a way to detect sample mishandling or other kinds of clerical error. However, in a data set of this size, some such mismatches would be expected due to transgender individuals, or instances of real (but rare) genetic variation, such as sex-chromosome aneuploidies [30]. Affymetrix genotype calling on the X and Y chromosomes allows only haploid or diploid genotype calls, depending on the inferred sex [29]. Therefore, cases of full or mosaic sex chromosome aneuploidies may result in compromised genotype calls on all, or parts of, the sex chromosomes (but not affect the autosomes). For example, individuals with karyotype XXY will likely have poorer quality genotype calls on the pseudoautosomal region (PAR) of the X chromosome, as they are effectively triploid in this region. Using information in the measured intensities of chromosomes X and Y, we identified a set of 652 (0.134%) individuals with sex chromosome karyotypes putatively different from XY or XX (**Figure 2B, Supplementary Table S1**). The list of samples is provided as part of the data release. Researchers wanting to identify sex mismatches should compare the self-reported sex and inferred sex data fields.

We did not remove samples from the data as a result of any of the above analyses, but rather provide the information as part of the data release. However, we excluded a small number of samples (835 in total) that we identified as sample duplicates (as opposed to identical twins, see **Supplementary Material**) or were likely involved in sample mishandling in the laboratory (~10), as well as participants who asked to be withdrawn from the project prior to the data release (33).

#### 2.1.6 Summary of genotype data quality

The application of our QC pipeline resulted in the released dataset of 488,377 samples and 805,426 markers from both arrays with the properties shown in **Figure 3**. The proportion of all the genotype calls made by Affymetrix that were set to missing as a result of quality control is 0.0097 (see **Table 3**). The proportion of genotyped samples that were identified as poor quality is 0.002 (968/488377). Furthermore, a set of 588 pairs of experimental duplicates show very high concordance. On average, 99.87% of a pair’s genotype calls are identical and the lowest rate is still 99.39% (**Supplementary Figure S13**).

**Figure 3.**
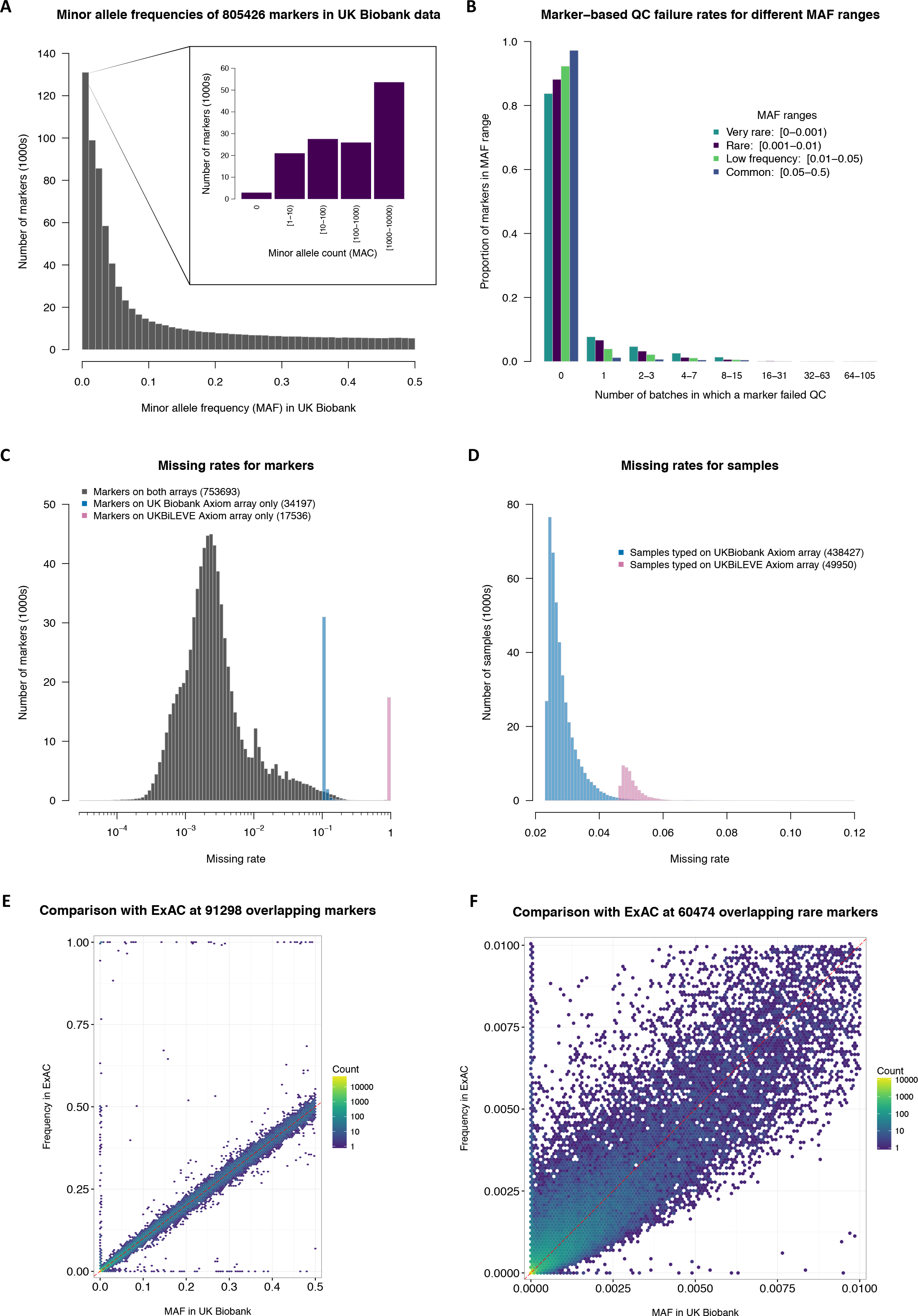
Summary of genotype data quality and content. All plots show properties of the UK Biobank genotype data that is made available to researchers, after applying QC as described in the main text. **(A)** Minor allele frequency (MAF) distribution. All samples were used in the calculation of MAF. The inset shows the number of markers within different minor allele count bins for rare markers only (MAF < 0.01). (**B)** The distribution of the number of batch-level QC tests that a marker fails (see **Table 2**; Methods). For each of four different MAF ranges (indicated by colours) we show the fraction of markers that fail the specified number of batches. For example, just over 92% of the ‘low frequency’ markers (0.01 ≤ MAF < 0.05) do not fail quality control in any batches. Any marker that failed all 106 batches is excluded from the data release, so such markers are not included here (see **Table 3. (C)** The distribution of missing rates for markers. Three histograms are overlaid, each showing a different, mutually exclusive, subset of markers, which are indicated by the three colours. Markers from only one array exhibit more missing data because only a subset of samples were typed on each array (10% on UK BiLEVE Axiom™ Array and 90% on the UK Biobank Axiom™ Array). **(D)** The distribution of missing rates for samples. Two histograms are overlaid, each showing a mutually exclusive subset of samples. All samples have some missing data because not all markers were included on both arrays (~2% are exclusive to the UK BiLEVE Axiom™ Array and ~4% exclusive to the UK Biobank Axiom™ Array). Additional missing data is also introduced from the batch-based marker QC. **(E)** Comparison of MAF in UK Biobank with the frequency of the same allele in ExAC, among the European-ancestry samples within each study. We used 33,370 samples for ExAC and 463,844 samples for UK Biobank (see **Supplementary Material** for details), and only markers that have a call rate greater than 0.9 in both studies. Each hexagonal bin is coloured according to the number of markers falling in that bin, as indicated by the key (note the log_10_ scale). The dashed red line shows x=y. The set of markers with very different allele frequencies seen on the top, bottom, and left-hand sides of the plot comprise ~300 markers. This is ~0.3% of all markers in the comparison, or ~0.5% of all markers with MAF > 0.001 in at least one study (see **Supplementary Material** for a discussion). **(F)** As with (E), but zooming in on the rare markers (both axes < 0.01).

Subsequent to the interim release of genotypes for ~150,000 UK Biobank participants improvements were made to the genotype calling algorithm [22] and quality control procedures. We therefore expect to observe some changes in the genotype calls and missing data profile of samples included in both the interim data release and this final data release. Discordance among non-missing markers is very low (mean 6.7x10^-5^; **Supplementary Figure S1**); and for each sample there are ~24,500 genotype calls (on average) that were missing in the interim data, but which have non-missing calls in this release. This is much smaller in the reverse direction, with ~500 calls, on average, missing in this release but not missing in the interim data, so there is an average net gain of ~24,000 genotype calls per sample.

We compared allele frequencies in the UK Biobank with those estimated from sequencing data sourced from the Exome Aggregation Consortium (ExAC) database [31]. We computed allele frequencies at a set of 91,298 overlapping markers, using samples with European ancestry. We do not expect allele frequencies in the two studies to match exactly due to subtle differences in the ancestral backgrounds of the individuals in each study, as well as differences in the sensitivity and specificity of the two technologies (exome sequencing and genotyping arrays). Despite this, overall the allele frequencies are encouragingly similar (r^2^ = 0.94) (**Figure 3E**). A discussion of the discrepancies between UK Biobank and ExAC is given in the **Supplementary Material**.

Over 110,000 rare markers (MAF < 0.01) were included on the two arrays used for the UK Biobank cohort [7]. Variants occurring at very low frequencies present a particular challenge for genotype calling using array technology, especially in cases where only a small number of samples within a genotyping batch are expected to have a copy of the minor allele (almost always with a heterozygous genotype)^2^. In these cases it is sometimes difficult in the genotype intensity data to distinguish a sample that genuinely has the minor allele, from one whose intensities are in the tails of the distribution of those in the major homozygote cluster. Examples of two different markers with MAF < 0.001 in UK Biobank are shown in **Figure 4B** and **Figure 4C**. One marker is performing well (4B), but for the other marker (4C) the heterozygous samples are more difficult to identify. In contrast, **Figure 4A** shows a common SNP (MAF=0.077), which has three well-separated clusters corresponding to the three genotypes. A larger fraction of rare markers fail quality control tests compared to low frequency and common markers, but 84% still pass in all batches (**Figure 3B**). The MAF of rare markers are generally similar to those in ExAC, but very rare markers tend to have a lower MAF in the UK Biobank (**Figure 3F**). We strongly recommend researchers visually inspect similar plots for markers of interest using a utility such as Evoker [32] (see **URLs**), especially for rare markers.

**Figure 4.**
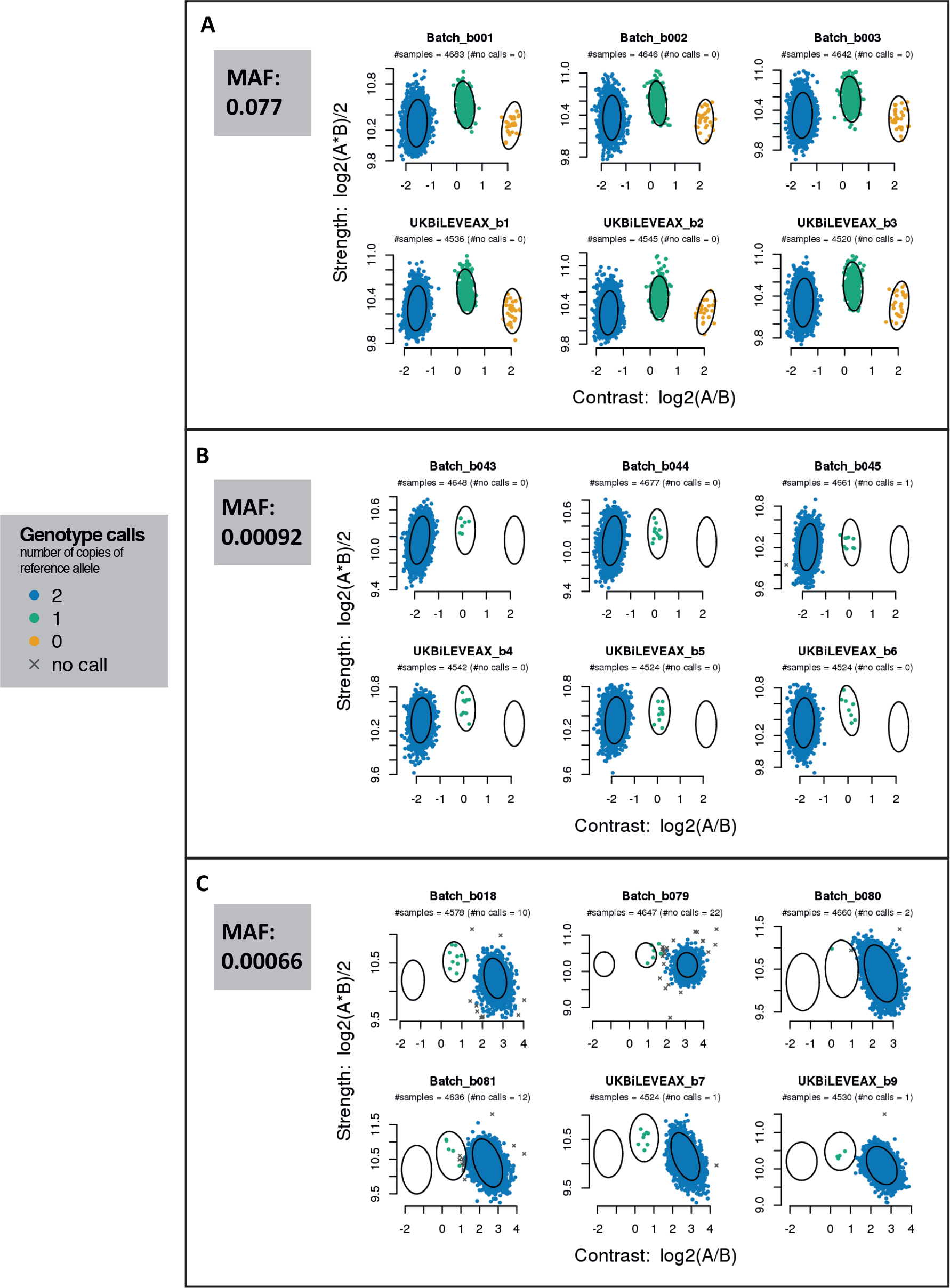
Examples of intensity data and genotype calls for markers of different allele frequencies. Each of the three sub-figures shows intensity data for a single marker within six different batches. Batches labelled with the prefix “UKBiLEVEAX” contain only samples typed using the UK BiLEVE Axiom array, and those with the prefix “Batch” contain only samples typed using the UK Biobank Axiom array. Each point represents one sample and is coloured according to the inferred genotype at the marker. The x and y axes are transformations of the intensities for probe sets targeting each of the alleles “A” and “B” (see **Supplementary Material** for definition of probe set). The ellipses indicate the location and shape of the posterior probability distribution (2-dimensional multivariate Normal) of the transformed intensities for the three genotypes in the stated batch. That is, each ellipse is drawn such that it contains 85% of the probability density. See [29] for more details of Affymetrix genotype calling. The minor allele frequency of each of the markers is computed using all samples in the released UK Biobank genotype data. **(A)** A marker with a MAF of 0.077 with well-separated genotype clusters. **(B)** Intensities for a marker with a MAF of 0.00092 with well-separated genotype clusters. As would be expected under HWE, there are no instances of samples with the minor homozygote genotype. **(C)** Intensities for a marker with a MAF of 0.00066, and where the heterozygote cluster is not well separated from the large major homozygote cluster in some batches, making it more difficult to confidently call the heterozygous genotypes.

In addition to the QC described above, a close examination of results of a test genome-wide association study (GWAS) (discussed in **Section 2.5**) provided further evidence for the quality of the resulting set of genotype calls.

### 2.2 Ancestral diversity and cryptic relatedness

#### 2.2.1 Genetic population structure

The diversity in ancestral origins of UK Biobank participants is evident from the self reported ethnic background (**Table 2**) and country of birth information. The genotype data provides a unique opportunity to study their ancestral origins in a quantitative manner. Accounting for the ancestral background of participants is an essential component of analysis of the UK Biobank resource, both for epidemiological studies [33], and genetic analyses, such as GWAS [27, 34]. We used principal components analysis (PCA) to measure population structure within the UK Biobank cohort. PCA is widely used as a method for assessing and potentially controlling for population structure in GWAS [27, 35].

We computed principal components (PCs) using an algorithm (*fastPCA* [36]) which performs well on datasets with hundreds of thousands of samples by approximating only the top *n* PCs that explain the most variation, where *n* is specified in advance. We computed the top 40 PCs using a set of 407,219 unrelated, high quality samples and 147,604 high quality markers pruned to minimise linkage disequilibrium (LD) [37]. We then computed the corresponding PC-loadings and projected all samples onto the PCs, thus forming a set of PC scores for all samples in the cohort (**Supplementary Material**).

**Figure 5** shows results for the first 6 PCs plotted in consecutive pairs. Results for further PCs are shown in **Supplementary Figures S5** and **S6**. As expected, individuals with similar PC scores have similar self-reported ethnic backgrounds. For example, the first two PCs separate out individuals with sub-Saharan African ancestry, European, and east-Asian ancestry. Individuals who self-report as mixed ethnicity tend to fall on a continuum between their constituent groups (e.g the White and Asian category). Further PCs capture population structure at sub-continental geographic scales. **Supplementary Figure S7** shows the relationship between each of the 40 PCs, and country of birth as reported by participants. For example, high scores in PC 6 are associated with individuals born in Central and South American countries such as Peru, Colombia, and Chile.

**Figure 5.**
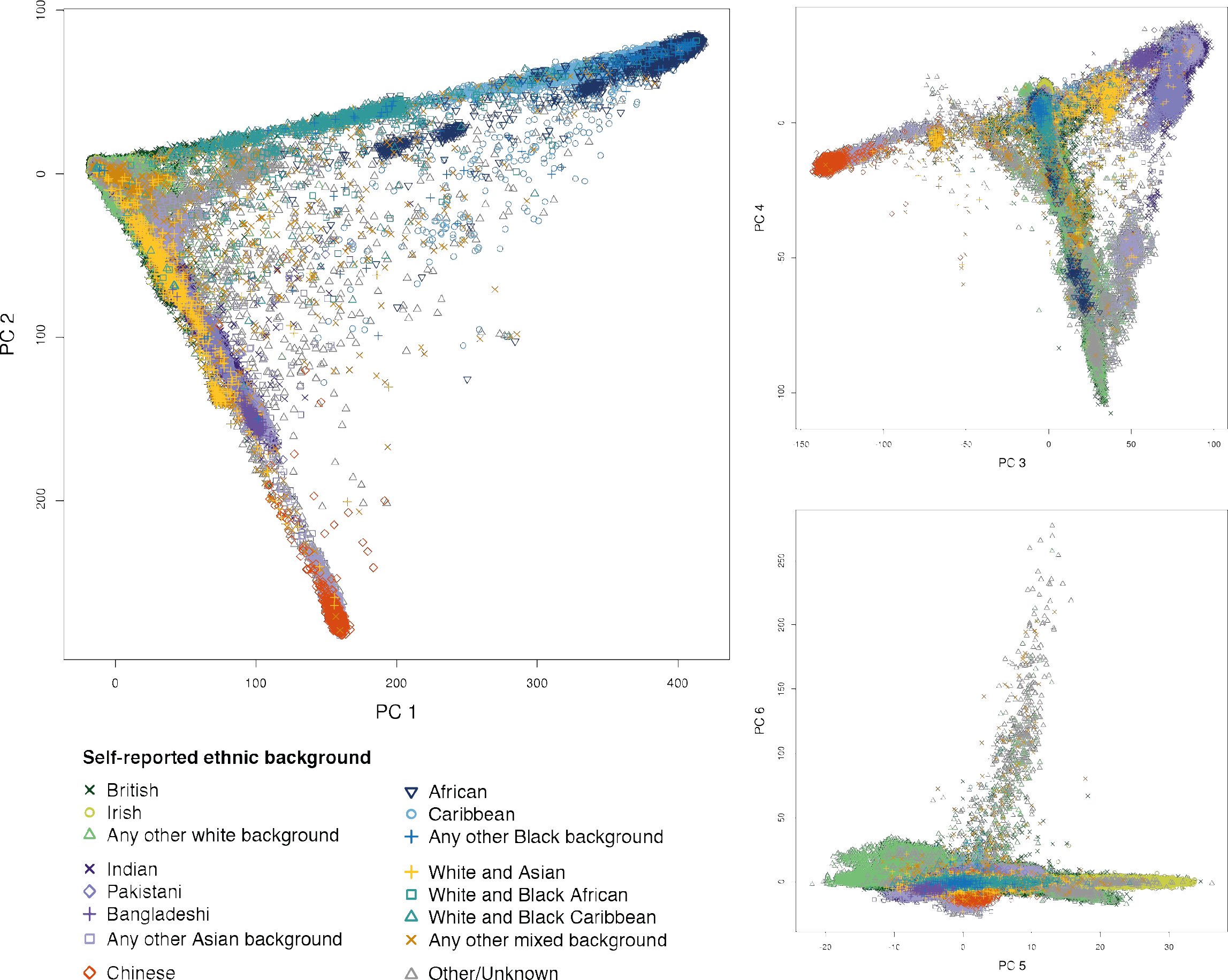
Ancestral diversity in the UK Biobank cohort. Plots of consecutive pairs of the first six principal components in a PCA of genotype data for UK Biobank participants (see **Supplementary Material**). Each point represents an individual and is placed according to their principal component scores (using genetic data only), with shapes and colours indicating their self-reported ethnic background as shown in the legend. See **Table 2** for the proportions of participants in each category. Plots of all pairs of these 6 PCs are shown in **Supplementary Figure S5**, and results of further PCs are represented in **Supplementary Figures S6** and **S7**.

#### 2.2.2 White British ancestry subset

Researchers may want to only analyse a set of individuals with relatively homogeneous ancestry to reduce the risk of confounding due to differences in ancestral background. Although the UK Biobank cohort is ethnically diverse, such analysis is feasible without compromising too much in sample size because a majority of participants in the UK Biobank cohort report their ethnic background as “British”, within the broader-level group “White” (88.26%). Our PCA revealed population structure even within this category (**Supplementary Figure S8**), so we used a combination of self-reported ethnic background and genetic information to identify a subset of 409,728 individuals (84%) who self-report as “British” and who have very similar ancestral backgrounds based on results of the PCA (see **Supplementary Material**). Fine-scale population structure is known to exist within the UK [38] but methods for detecting such subtle structure [39] available at the time of analysis are not feasible to apply at the scale of the UK Biobank. The white British ancestry subset may therefore still contain subtle structure present at subnational scales.

#### 2.2.3 Cryptic relatedness

Close relationships (e.g siblings) among UK Biobank participants were not recorded during the collection of other phenotypic information. Indeed, many participants may not be aware that a close relative (such as an aunt, or sibling) is also part of the cohort. This information can be important for epidemiological analyses [40], as well as in GWAS [41], and the genetic data provides an opportunity to discover and characterise familial relatedness within the cohort. This analysis, combined with phenotype information, is also useful for identifying samples that are experimental duplicates rather than genuine twins (see **Supplementary Material**).

We identified related individuals by estimating kinship coefficients for all pairs of samples, and report coefficients for pairs of relatives who we infer to be 3^rd^ degree or closer. We used an estimator implemented in the software, *KING* [42], as it is robust to population structure (i.e does not rely on accurate estimates of population allele frequencies) and it is implemented in an algorithm efficient enough to consider all pairs (~1.2x10^11^) in a practicable amount of time. As noted by the authors of *KING* [42], we found that recent admixture (e.g “Mixed” ancestral backgrounds) tended to inflate the estimate of the kinship coefficient, as the estimator assumes HWE among markers with the same underlying allele frequencies within an individual. We alleviated this effect by only using a subset of markers that are only weakly informative of ancestral background (see **Supplementary Material, Figure S12**). We also excluded a small fraction of individuals (977) from the kinship estimation, as they had properties (e.g high missing rates) that would lead to unreliable kinship estimates (see **Supplementary Material**). We called relationship classes for each related pair using the kinship coefficient and fraction of markers for which they share no alleles (IBS0), and using the boundaries recommend by the authors of *KING*.

A total of 147,731 UK Biobank participants (30.3%) are inferred to be related (3^rd^ degree or closer) to at least one other person in the cohort, and form a total of 107,162 related pairs (**Table 4**). This is a surprisingly large number, and it is not driven solely by an excess of 3^rd^ degree relatives. For example, the number of sibling pairs (22,666) is about twice as many as would theoretically be expected in a random sample (of this size) of the eligible UK population, after taking into account typical family sizes (see **Supplementary Material**). To ensure we were not overestimating the number of related pairs, we inferred related pairs (within a subset of the data) using a different inference method implemented in *PLINK* (“--genome” command) [43] and confirmed 100% of the twins, parent-offspring and sibling pairs, and 99.9% of pairs overall (see **Supplementary Material**). The larger than expected number of related pairs could be explained by sampling bias due to, for example, an individual being more likely to agree to participate because a family member was also involved. Furthermore if, as seems plausible, related individuals cluster geographically, rather than being randomly located across the United Kingdom (UK), the assessment-centre based recruitment strategy employed by UK Biobank [1] will naturally tend to oversample related individuals, relative to random sampling of the UK population.

**Table 4.** Summary of related pairs (3rd degree or closer) for the full UK Biobank cohort. Counts are derived from the kinship coefficients as recommended by the authors of KING [42]. Note that parent-offspring and full sibling pairs have the same expected kinship coefficient (0.25) but can be easily distinguished by their IBS0 fraction. The count of monozygotic twins is after excluding samples identified as duplicates (see Supplementary Material).

|  | Monozygotic twins | Parent-offspring | Full siblings | 2<sup>nd</sup> degree | 3<sup>rd</sup> degree | Total |
| --- | --- | --- | --- | --- | --- | --- |
| Number of pairs | 179 | 6,276 | 22,666 | 11,113 | 66,928 | 107,162 |

#### 2.2.4 Trios and extended families

Pairs of related individuals within the UK Biobank cohort form networks of related individuals, or ‘family’ groups. In most cases these are of size two, but there are many groups of size 3 or larger in the cohort, even when restricting to 2^nd^ degree or closer relative pairs. By considering the relationship types and the age and sex of individuals within each family group, we identified 1,066 sets of trios (two parents and an offspring), which comprises 1,029 unique sets of parents and 37 quartets (two parents and two children). There are no instances of 3-generation nuclear families (grandparent-parent-offspring), which is not surprising, given that the age-range of the cohort spans only 30 years.

There are 172 family groups with 5 or more individuals that are related to 2^nd^ degree or closer, and **Figure 6** illustrates examples of large nuclear families in the cohort. One of these is a group of eleven individuals where every pair is a 2^nd^ degree relatives. The simplest explanation for such a family group is that all of the individuals are half siblings, with one shared parent who is not in the cohort. Alternatively, one individual in the group could be a sibling or a parent of the shared parent. The pattern of haplotype sharing between the individuals would distinguish these cases but we have not undertaken this analysis. We did, however, confirm that the shared parent must be their father, because the individuals do not all carry the same Mitochondrial alleles, and all the males in the group have the same alleles on their Y chromosome (data not shown).

**Figure 6.**
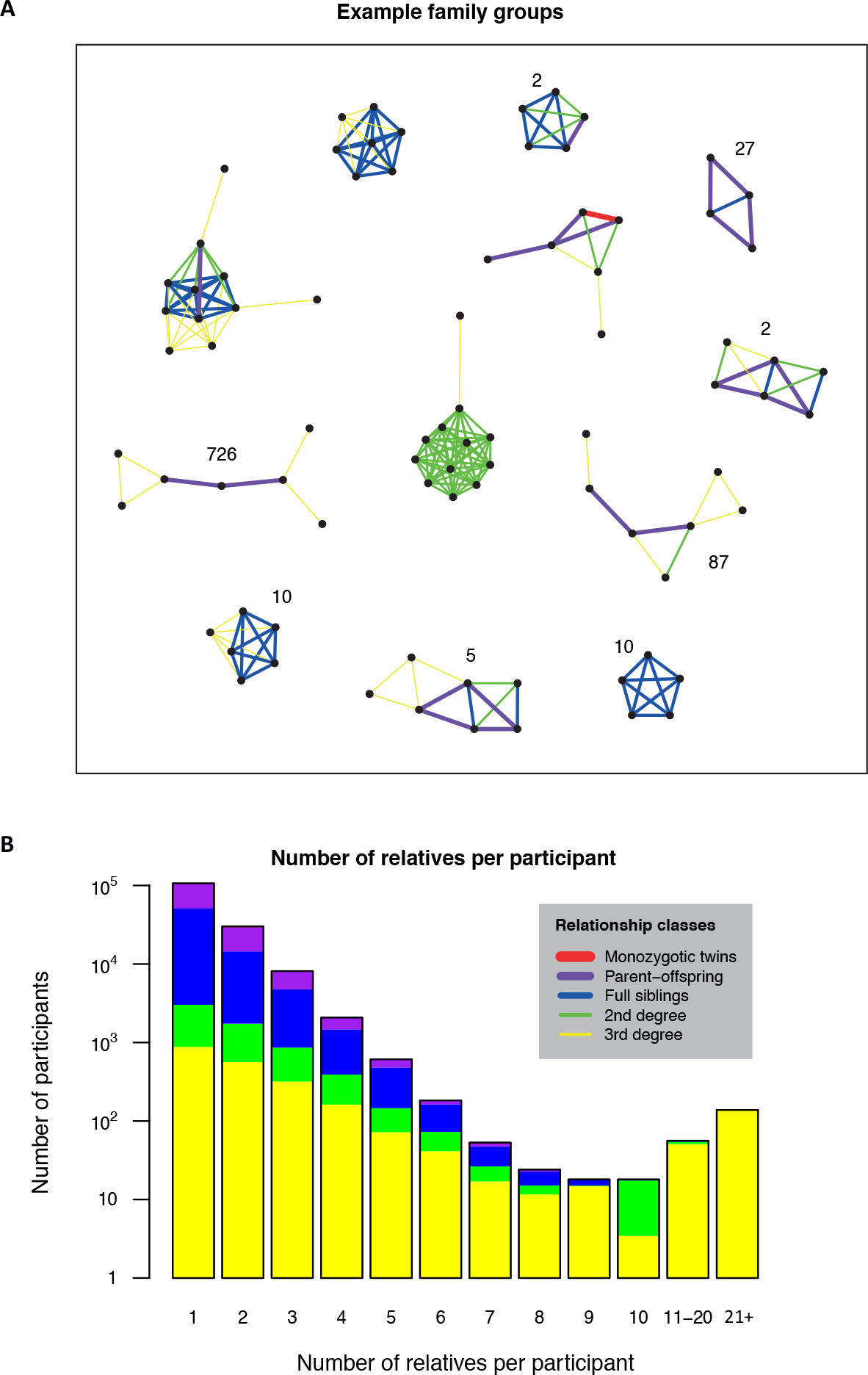
Familial relatedness in the UK Biobank cohort. (A) Examples of family groups within the UK Biobank cohort. Points indicate participants, and lines between points indicate familial relatedness (3^rd^ degree and closer) as inferred from the genetic data (see Methods). The colour and thickness of the lines indicate different relative classes, as shown in the key. An integer next to a network indicates the total number of family networks in the cohort with the same configuration, ignoring 3^rd^ degree pairs. No integer means there is only the one shown. For example, there are 10 networks that comprise exactly 5 full siblings (two examples, which differ with respect to a 3^rd^ degree relative, are shown on this plot); and there is only one network that comprises 6 full siblings (plus one 3^rd^ degree relative who is related to all siblings). **(B)** Distribution of the exact number of relatives that participants have in the UK Biobank cohort. The height of each bar shows the count of participants who have exactly the stated number of relatives. Note the logarithmic scale. The colours indicate the proportions of each relatedness class for individuals counted within a bar. For example, for each individual that has three relatives in the cohort (3^rd^ bar), we count how many of their relatives are in each relatedness class (i.e a full sibling, parent-offspring etc.). We then sum these counts over all individuals with three relatives and colour the bar according to the proportions of each relatedness class. In this group ~20% of their relatives are full siblings and ~64% are 3^rd^ degree relatives. There are also 18 participants with exactly 10 relatives. The unusually large fraction of 2^nd^ degree relationships for this group is a result of the set of eleven individuals who are all 2^nd^ degree relatives of each other, as shown in the centre of (A).

### 2.3 Phasing and Imputation of SNPs, short indels and CNVs

Genotype imputation [44] is the process of predicting genotypes that are not directly assayed in a sample of individuals. A reference panel of haplotypes at a dense set of SNPs, indels and structural variants (SVs), is used to impute genotypes into a study sample of individuals that have been genotyped at a subset of the markers. These ‘*in silico*’ genotypes can then be used to boost the number of markers that can be tested for association. This increases the power of the study, the ability to resolve or fine-map the causal variants and facilitates meta-analysis. The process of imputation first involes pre-phasing the directly genotyped markers, followed by a haploid imputation step [45].

#### 2.3.1 Pre-Phasing

For the pre-phasing step we only used markers present on both the UK BiLEVE and UK Biobank Axiom arrays. We removed markers which (a) failed QC in > 1 batch, (b) had more than 5% missingness (c) had a minor allele frequency < 0.0001. We removed samples that were identified as outliers for heterozygosity and missingness (**Section 2.1.5**). These filters resulted in a dataset with 670,739 autosomal markers in 487,442 samples. Phasing on the autosomes was carried out using *SHAPEIT3* [46] in chunks of 15,000 markers, with an overlap of 250 markers between chunks. Each chunk used 4 cores per job and S=200 copying states. The 1000 Genomes Phase 3 dataset [47] was used as a reference panel, predominantly to help with the phasing of samples with non-European ancestry. Chunks were ligated using a modified version of the *hapfuse* program (see **URLs**).

We assessed the accuracy of the phasing in a separate experiment by taking advantage of mother-father-child trios that were identified in the UK Biobank cohort (see **Section 2.2.4**). This family information can be used to infer the phase of a large number of markers in the trio parents. These family-inferred haplotypes were used as a truth set, as is common in the phasing literature. The parents of each trio were removed from the dataset and then haplotypes were estimated across chromosome 20 in a single run of *SHAPEIT3*. This dataset consisted of 16,175 autosomal markers.

The inferred haplotypes were then compared to the truth set using the switch error metric. Using a set of 696 trios with self-reported ethnic background “British” (within the broader-level group “White” (see **Table 2**)) and no other twins or 1^st^ or 2^nd^ degree relatives in the UK Biobank dataset, we estimated a median switch error rate of 0.229%. We also used a subset of 397 of these trios that also had no 3^rd^ degree relatives and obtained a median switch error rate of 0.234%.

#### 2.3.2 Reference panel used for imputation

There are a number of factors that influence the accuracy of genotype imputation, but generally accuracy will increase as the number of haplotypes in the reference panel grows and if the ancestry of the sample haplotypes is a good match to the ancestry of the reference panel haplotypes. The UK Biobank dataset consists of samples with a diverse range of ancestries, but with the majority of samples having British (or European) ancestry (**Table 2**). For this reason it was desirable to use a reference panel with a large number of haplotypes with British and European ancestry, and also a diverse set of haplotypes from other world-wide populations.

For the interim data release a reference panel was used that merged the UK10K and 1000 Genomes Phase 3 reference panels, which consisted of 87,696,888 bi-allelic markers in 12,570 haplotypes. An advantage of this reference panel is that it includes short indels and larger structural variants as well as SNPs. Since then, the HRC reference panel [16] has been widely adopted for imputation into many GWAS. This reference panel has many more haplotypes (64,976) than the previously used reference panel, and so is expected to produce better imputation performance, especially at lower allele frequencies (see **Supplementary Figure S14**). However the HRC panel has fewer SNPs (39,235,157) since very rare variants were excluded when the panel was constructed, and there are no short indels and structural variants. To obtain the expected gain in performance at HRC SNPs, whilst retaining results at all the SNPs/indels/SVs imputed in the interim release, we imputed SNPs from both panels, but preferentially used the HRC imputation at SNPs present in both panels.

#### 2.3.3 Imputation

To facilitate fast imputation of all ~500,000 samples we re-coded *IMPUTE2* [45] to focus exclusively on the haploid imputation needed when samples have been pre-phased. This new version of the program is referred to as *IMPUTE4* (see **URLs**), but uses exactly the same hidden Markov model (HMM) within *IMPUTE2*, and produces identical results to *IMPUTE2* when run using all reference haplotypes as hidden states (data not shown). To reduce RAM usage and increase speed we used compact binary data structures and took advantage of high correlations between inferred copying states in the HMM to reduce computation. Imputation was carried out in chunks of approximately 50,000 imputed markers with a 250kb buffer region and on 5,000 samples per compute job. The combined processing time per sample for the whole genome was ~10mins.

The result of the imputation process is a dataset with 92,693,895 autosomal SNPs, short indels and large structural variants in 487,442 individuals. dbSNP Reference SNP (rs) IDs were assigned to as many markers as possible using rs ID lists available from the UCSC genome annotation database for the GRCh37 assembly of the human genome (see **URLs**).

**Figure 7** shows the distribution of information scores on all markers in the imputed dataset. An information score of α in a sample of *M* individuals indicates that the amount of data at the imputed marker is approximately equivalent to a set of perfectly observed genotype data in a sample size of α*M*. The figure illustrates that the majority of markers above 0.1% frequency have high information scores. Previous GWAS have tended to use a filter on information around 0.3 which roughly corresponds to an effective sample size of ~150,000. Thus it maybe possible to reduce the information score threshold and still obtain good power to detect associations.

**Figure 7.**
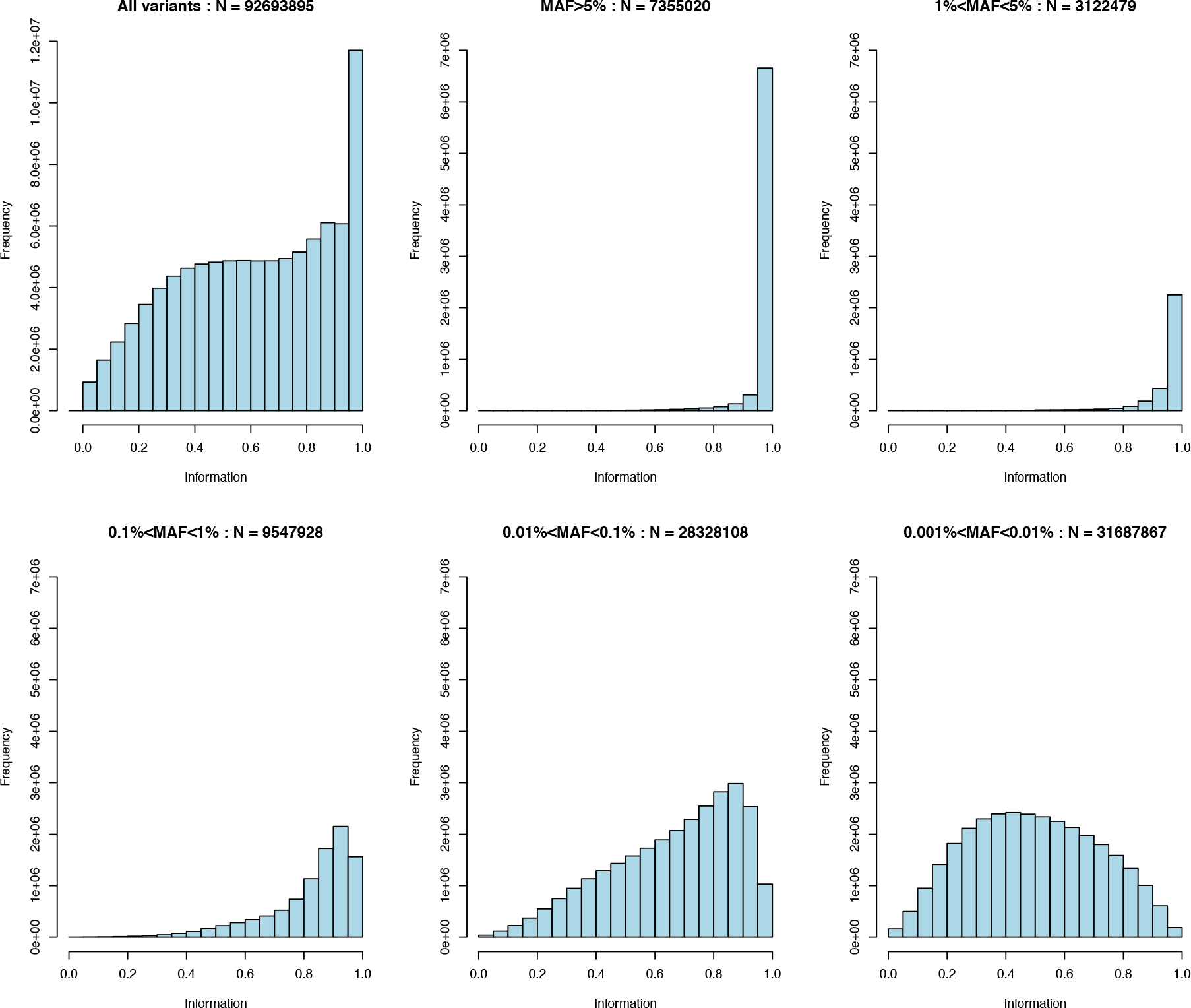
Distribution of information scores at autosomal markers in the imputed dataset. Top left shows the full distribution of the info scores. The remaining panels show distributions in tranches of minor allele frequency MAF > 5%, 1%<=MAF<5%, 0.1%<=MAF<1%, 0.01%<=MAF<0.1% and 0.001%<=MAF<0.01%.

The UK Biobank Axiom array from Affymetrix was specifically designed to optimize imputation performance in GWAS studies [7]. An experiment was carried out to assess the imputation performance of the array, stratified by allele frequency, and to compare performance to a range of old and new commercially available arrays (**Supplementary Material** and **Figure S14**). These results highlight the very good performance of the Axiom array compared to other arrays across the full frequency spectrum.

#### 2.3.4 A compressed and indexed data format for imputed data

The interim release of the UK Biobank genetic data was made available in version 1.1 of the BGEN file format, which is a binary version of the GEN file format (see **URLs**). The interim data release imputed files containing ~150,000 samples at ~73M autosomal SNPs require 1.3Tb of file space.

For the full data release we developed a new version (version 1.2) of the BGEN file format, providing greater compression and the ability to store phased haplotype data. The full imputed files containing ~500,000 samples at ~93M autosomal markers require 2.1Tb of file space. In addition, we developed an open-source software library and suite of tools that can be used to manipulate and access the BGEN files, including the tool *BGENIX*, which provides random access to data by making use of a separate index file (see **URLs**).

Several commonly-used programs support the BGEN format, including *SNPTEST* (see **URLs**) and *PLINK* [48]. The program *QCTOOL v2* (see **URLs**) can be used to filter, summarize, manipulate and convert the files to other formats.

### 2.4 Imputation of classical HLA alleles

The major histocompatibility complex (MHC) on chromosome six is the most polymorphic region of the human genome and harbours the largest number of genetic associations to common diseases [49]. The relevance of this genomic region to biomedical research means that high quality typing of HLA alleles is a valuable extension to the genetic information in the UK Biobank. Because of the extensive genetic variation and strong linkage-disequilibrium in the MHC we used a specialised algorithm to resolve allelic variation at 11 classical human leukocyte antigen (HLA) genes in the UK Biobank cohort. We imputed HLA types at two-field (also known as four-digit) resolution for *HLA-A, -B, -C, -DRB5, -DRB4, -DRB3, -DRB1, -DQB1, -DQA1, - DPB1* and *-DPA1*, using the HLA*IMP:02 algorithm with a multi-population reference panel (**Supplementary Tables S4, S5**) [50, 51].

We estimated the accuracy of the imputation process using five-fold cross-validation in the reference panel samples with markers common to the reference panel and UK Biobank genotype data. For samples of European ancestry, the estimated four-digit accuracy for the maximum posterior probability genotype is above 93.9% for all 11 loci (**Supplementary Table S6**). This accuracy improved to above 96.1% for all 11 loci after restricting to HLA allele calls with a posterior probability greater than 0.70. This resulted in call rates above 95.1% for all loci (**Supplementary Table S7**).

To demonstrate the utility of the HLA imputation we performed association tests for diseases known to have HLA associations. We analysed 409,724 individuals in the white British ancestry subset (defined in Section 2.2.2) along with 446 disease codes based on self-reported diagnosis terms. Of these disease codes we focused on 11 immune-mediated diseases with known HLA associations (see **Supplementary Table S8** for a full list). For each individual we defined the HLA genotype at each locus as the pair of alleles with maximum posterior probability. We performed a standard association analysis (see for example [52]) for HLA alleles and each disease using logistic regression with an additive model for each allele at each locus. This effectively treats each allele as a SNP in a standard GWAS framework. For further details see the **Supplementary Material Section S5**. To ensure we only include high quality HLA calls in our analyses we performed a similar analysis where at each locus we only include those individuals whose genotype pair has a maximum posterior probability greater than 0.7. No significant differences were observed compared to the full analysis (data not shown). For each disease in our analysis we identified the HLA allele with the strongest evidence of association and these were consistent with previous reports (see **Supplementary Table S8** for the full results). This provides evidence that the imputation of HLA alleles is sufficiently accurate for genetic association testing.

As a final QC check for our imputed HLA alleles, we show that the UK Biobank data provides consistent results with a fine-mapping study. In this case we replicate independent HLA associations in a single disease study of multiple sclerosis susceptibility by the International Multiple Sclerosis Genetics Consortium (IMSGC) [52]. Here we observed evidence of association and effect size estimates for HLA alleles that are consistent with those found in IMSGC study (**Table 5**).

**Table 5.**
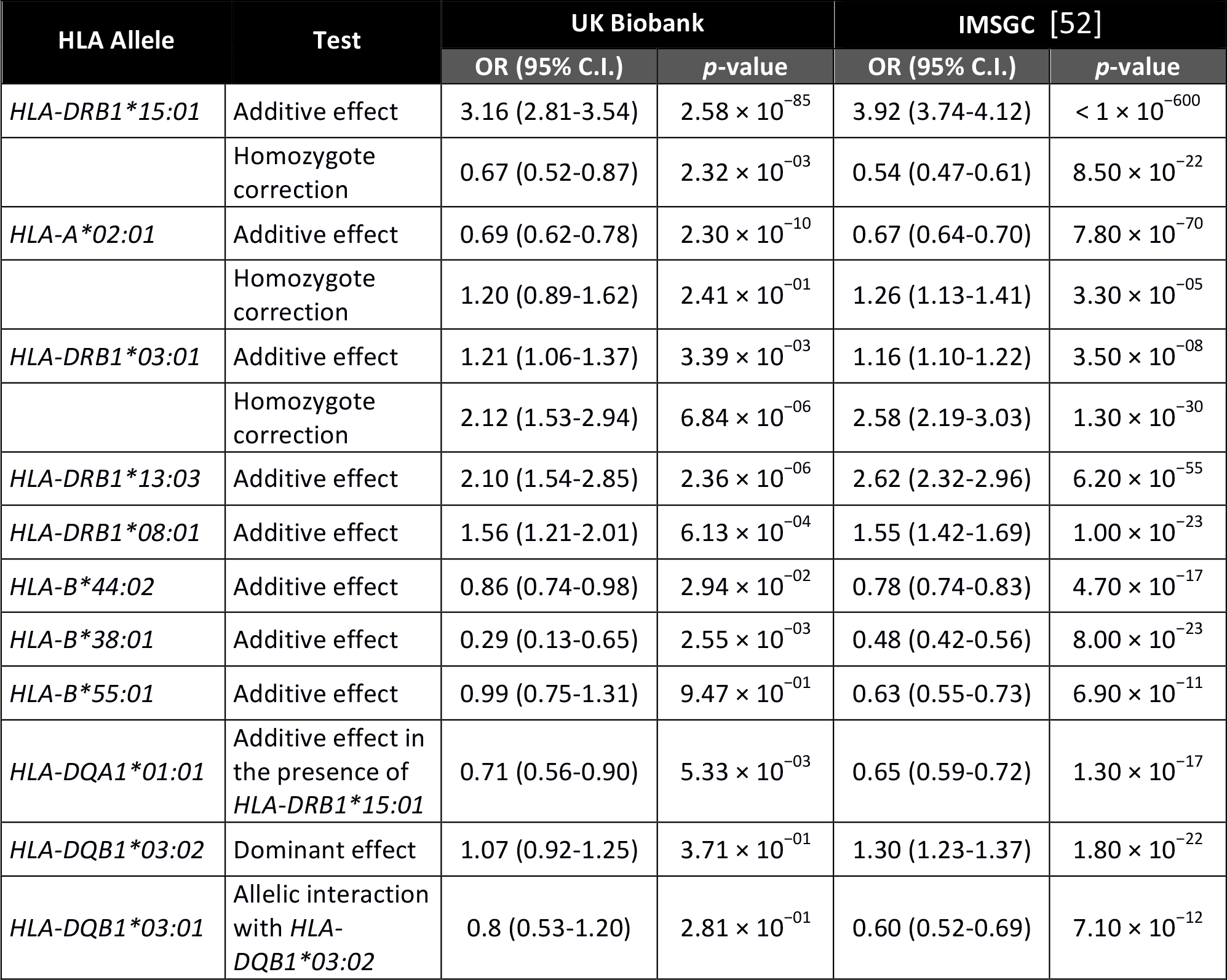
Evidence of association between HLA alleles and multiple sclerosis in UK Biobank compared to the IMSGC cohort. The UK Biobank association tests involved 1,501 self-reported cases and 409,724 controls; the IMSGC cohort involved 17,465 cases and 30,385 controls [52]. Thus the UK Biobank analysis had significantly lower power than the IMSGC analysis, which is reflected in the reported *p*-values. Effect sizes for the UK Biobank were estimated jointly using the model of the MHC reported by the IMSGC (with the exception of the two SNPs rs9277565 and rs2229029). As in the IMSGC analysis the homozygote correction test indicates a departure from additivity. That is, if the odds ratio is < 1 then the homozygous effect is smaller than under the additivity assumption and bigger if it is > 1.

| HLA Allele | Test | UK Biobank |  | IMSGC [52] |  |
| --- | --- | --- | --- | --- | --- |
|  |  | OR (95% C.I.) | p-value | OR (95% C.I.) | p-value |
| *HLA-DRB1\*15:01* | Additive effect | 3.16 (2.81-3.54) | $2.58 \times 10^{-85}$ | 3.92 (3.74-4.12) | $< 1 \times 10^{-600}$ |
| | Homozygote correction | 0.67 (0.52-0.87) | $2.32 \times 10^{-03}$ | 0.54 (0.47-0.61) | $8.50 \times 10^{-22}$ |
| *HLA-A\*02:01* | Additive effect | 0.69 (0.62-0.78) | $2.30 \times 10^{-10}$ | 0.67 (0.64-0.70) | $7.80 \times 10^{-70}$ |
| | Homozygote correction | 1.20 (0.89-1.62) | $2.41 \times 10^{-01}$ | 1.26 (1.13-1.41) | $3.30 \times 10^{-05}$ |
| *HLA-DRB1\*03:01* | Additive effect | 1.21 (1.06-1.37) | $3.39 \times 10^{-03}$ | 1.16 (1.10-1.22) | $3.50 \times 10^{-08}$ |
| | Homozygote correction | 2.12 (1.53-2.94) | $6.84 \times 10^{-06}$ | 2.58 (2.19-3.03) | $1.30 \times 10^{-30}$ |
| *HLA-DRB1\*13:03* | Additive effect | 2.10 (1.54-2.85) | $2.36 \times 10^{-06}$ | 2.62 (2.32-2.96) | $6.20 \times 10^{-55}$ |
| *HLA-DRB1\*08:01* | Additive effect | 1.56 (1.21-2.01) | $6.13 \times 10^{-04}$ | 1.55 (1.42-1.69) | $1.00 \times 10^{-23}$ |
| *HLA-B\*44:02* | Additive effect | 0.86 (0.74-0.98) | $2.94 \times 10^{-02}$ | 0.78 (0.74-0.83) | $4.70 \times 10^{-17}$ |
| *HLA-B\*38:01* | Additive effect | 0.29 (0.13-0.65) | $2.55 \times 10^{-03}$ | 0.48 (0.42-0.56) | $8.00 \times 10^{-23}$ |
| *HLA-B\*55:01* | Additive effect | 0.99 (0.75-1.31) | $9.47 \times 10^{-01}$ | 0.63 (0.55-0.73) | $6.90 \times 10^{-11}$ |
| *HLA-DQA1\*01:01* | Additive effect in the presence of *HLA-DRB1\*15:01* | 0.71 (0.56-0.90) | $5.33 \times 10^{-03}$ | 0.65 (0.59-0.72) | $1.30 \times 10^{-17}$ |
| *HLA-DQB1\*03:02* | Dominant effect | 1.07 (0.92-1.25) | $3.71 \times 10^{-01}$ | 1.30 (1.23-1.37) | $1.80 \times 10^{-22}$ |
| *HLA-DQB1\*03:01* | Allelic interaction with *HLA-DQB1\*03:02* | 0.8 (0.53-1.20) | $2.81 \times 10^{-01}$ | 0.60 (0.52-0.69) | $7.10 \times 10^{-12}$ |

### 2.5 GWAS for standing height

As a final demonstration of the quality of the directly genotyped and imputed data we conducted a genome-wide association scan for a well-studied [49], and highly polygenic human trait: standing height. Previously published, large cohort studies for this trait also provide a suitable independent comparison set for our scan. We conducted the scan using genotypes for a set of 343,321 unrelated UK Biobank participants within the white British ancestry subset (Section 2.2.2) and with a standing height measurement (**Supplementary Material**). We compared our results to the largest published GWAS for human height that does not use UK Biobank data: a meta analysis involving a total of 253,288 individuals of European ancestry carried out by the Genetic Investigation of Anthropometric Traits (GIANT) Consortium [53] in 2014.

**Figure 8** shows the p-values for associations with height on one chromosome, along with *p*-values for GIANT’s meta analysis. Reassuringly, the pattern of associations is very similar in both the UK Biobank and GIANT results. The gain in power in the UK Biobank cohort is clear, with many loci reaching genome-wide significance (*p*-value < 5x10^-8^) in the UK Biobank, which do not in the GIANT study (**Supplementary Figure S15**). The purpose of this analysis is not to report novel associations for height, rather to indicate the potential of the resource to uncover such findings. However, we chose to focus on a single associated region on chromosome 2 shown in **Figure 8D**. Correlations (r^2^) between markers in this region show a pattern that is as expected in the context of linkage disequilibrium (LD), and the local recombination rates. The stripe-like pattern of the association statistics is indicative of multiple mutations occurring on similar branches of the genealogical tree underlying the data, which are likely linked to varying degrees with the causal marker(s). The correlation (r^2^) between the most associated marker and all other markers in the region drops of sharply around the small peak in recombination (Hapmap recombination map [21]) to the right of the most significantly associated marker. Interestingly, this marker was imputed from the genotypes, which points to the success of the imputation in this study, and in general, to the value of imputing millions more markers.

**Figure 8.**
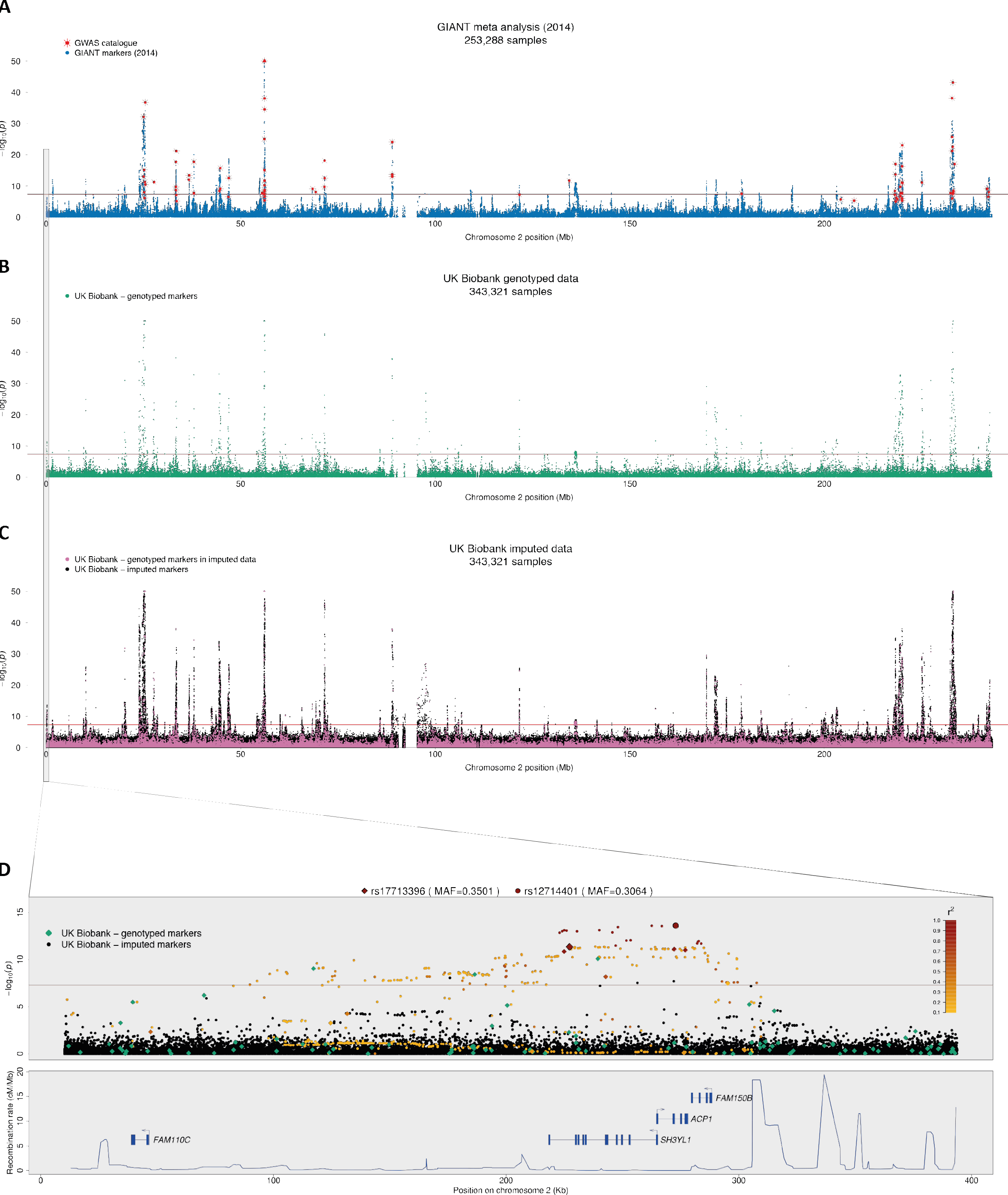
Association statistics for human height. Results (*p*-values) of association tests between human height and genotypes using three different sets of data for chromosome 2. *p*-values are shown on the –log_10_ scale and capped at 50 for visual clarity. Markers with –log_10_(*p*) > 50 are plotted at 50 on the y-axis and shown as triangles rather than dots. **(A)** Results for meta-analysis by GIANT (2014), with NCBI GWAS catalogue markers superimposed in red (plotted at the reported *p*-values). **(B)** Association statistics for UK Biobank markers in the genotype data. **(C)** Association statistics for UK Biobank markers in the imputed data. Points coloured pink indicate genotyped markers that were used in pre-phasing and imputation (see Section 2.3.1). This means that most of the data at each of these markers comes from the genotyping assay. Black points (the vast majority, ~8M) indicate fully imputed markers. **(D)** Results from all three data sets focussing on a ~3 Mega-base region at the terminal end of the p-arm. Genotyped markers (i.e markers in B) are shown as diamonds, and imputed markers (i.e. only markers coloured black in C) as circles. The two markers with the smallest *p*-value for each of the genotyped data and imputed data are enlarged and highlighted with black outlines, and other UK Biobank markers are coloured according to their correlation (r^2^) with one of these two. That is, genotyped markers with the leading genotyped marker (rs17713396), and imputed markers with the leading imputed marker (rs12714401). Markers with r^2^ less than 0.1 are shown as black or green.

Human height is a highly polygenic trait [53] and this provided an opportunity to examine many such regions of association. Other regions that we visually examined showed similar patterns, and all regions containing the variants reported to be associated with height in the NCBI GWAS catalogue (as of Feb 2017, excluding results based on UK Biobank data) were also genome-wide significant, or close to, in the UK Biobank data.

### 2.6 Multiple trait GWAS and PheWAS

To facilitate the running of GWAS for multiple continuous traits and fast PheWAS we have provided a new tool called *BGENIE* that is built upon the *BGEN* library (see **URLs**). The program also uses the *Eigen* matrix library and *OpenMP* to carryout as many of the linear algebra operations in parallel as possible. For example, estimation of effect sizes of large numbers of markers can be carried out in parallel using matrix operations, and we use indexing of missing data values to allow for fast estimation of standard errors. The program takes BGEN files as input and avoids repeated decompression and conversion of these files when analysing multiple phenotypes, and can lead to considerable time savings when compared to analysis using *PLINK* (see **Supplementary Material**).

## 3 Data provision and access

Genotype calls, both imputed and directly assayed, and HLA haplotype calls are available on a cost-recovery basis to researchers on successful application to the UK Biobank http://www.ukbiobank.ac.uk/scientists-3/genetic-data/. Several types of important information are also available along with the genotype calls, namely:

- Various quality control metrics and flags for samples and markers.
- 40 principal components for all genotyped samples, and kinship coefficients for pairs of inferred close relatives.
- Measured A and B allele intensities, log2 ratios and B-allele frequency data for all 488,377 genotyped samples at 805,426 markers (Affymetrix).
- Confidence values for all genotype calls in the released data.
- Posterior parameters for genotype clusters at each marker in each genotyping batch (Affymetrix).
- Imputation info scores and minor allele frequencies for all markers in the imputed data files.

## 4 URLs

*SHAPEIT3*, *IMPUTE4*, *BGENIE* - https://jmarchini.org/software/ *Hapfuse* - https://bitbucket.org/wkretzsch/hapfuse/src

*BGENIX, BGEN* library - https://bitbucket.org/gavinband/bgen GRCh37 human genome assembly - http://hgdownload.cse.ucsc.edu/goldenpath/hg19/database/ *Evoker* - https://github.com/wtsi-medical-genomics/evoker

BGEN file format - http://www.well.ox.ac.uk/~gav/bgen_format/bgen_format.html

GEN file format - http://www.stats.ox.ac.uk/~marchini/software/gwas/file_format.html

*SNPTEST* - https://mathgen.stats.ox.ac.uk/genetics_software/snptest/snptest.html

*QCTOOL v2* - http://www.well.ox.ac.uk/~gav/qctool_v2

## 5 Author contributions

C.B., C.F., D.P. carried out the marker and sample quality control and analysis. S.W. contributed to the genotype quality control. A.C., A.M., D.V., S.L., G.M. carried out all analysis relating to the HLA imputation and association testing. G.B. developed the BGEN file format. O.D., J.O., K.S., L.E. and J.M. carried out all analysis and methods development relating to the phasing, imputation and multiple trait analysis. C.B., C.F. and J.M. carried out all analysis relating to GWAS testing. J.M. and P.D. supervised the work. C.B., C.F., A.C., G.M., J.M., P.D. wrote the paper.

## Acknowledgements

### 6 Acknowledgements

This work was supported by the Wellcome Trust Core Awards 090532/Z/09/Z and 203141/Z/16/Z, and the European Research Council (ERC; grant 617306) (to J.M.) and Wellcome Trust Senior Investigator Awards 095552/Z/11/Z (to P.D.) and 100956/Z/13/Z (to G.M.), and UK Biobank (to C.B., D.P. and C.F.), and Wellcome Trust grant 100308/Z/12/Z (to A.C.), and Wellcome Trust grant 090770/Z/09/Z (to G.B.), and the Australian National Health and Medical Research Council (NHMRC), Career Development Fellowship ID 1053756 (to S.L.). The sample processing and genotyping was supported by the National Institute for Health Research, Medical Research Council, and British Heart Foundation.

We would like to acknowledge the Research Computing Core at the Wellcome Trust Centre for Human Genetics for their help in managing the huge computational workload of this project. We would like to acknowledge Affymetrix for their contribution to this project and specifically for their contribution to discussions of QC. We thank Alex Young for assistance with aspects of the relatedness analysis. We thank Alex Dilthey and Loukas Moutsianas for their assistance with performing classical HLA allele imputation and analysis. We would like to acknowledge UK Biobank co-ordinating centre staff for their role in extracting the DNA for this project. We would also like to express our gratitude to the participants in the UK Biobank cohort.

## 7 Conflicts of Interest

J.M. is a founder and director of Gensci Ltd. P.D., G.M. and S.L. are partners in Peptide Groove LLP. G.M. and P.D. are founders and directors of Genomics Plc in which P.D. currently holds an executive role. The remaining authors declare no competing financial interests.

## Footnotes

1 Now part of Thermo Fisher Scientific.

2 The genotyping batches in the UK Biobank project contain ~4,700 samples, so a MAF of, for example, 0.001 corresponds to an expected count (under HWE) of about 9 heterozygotes per batch.

